# Effect of cortisol on cortical organoids: Building a “stress in a dish” model system

**DOI:** 10.1101/2025.09.16.676476

**Authors:** Carolin Purmann, Kaela Farrise, Yiling Huang, Reenal Pattni, Marcus Ho, Victor G. Carrion, Alexander E. Urban

## Abstract

Exposure to chronic stress and traumatic experiences impacts brain health and development, which may lead to Post Traumatic Stress Disorder (PTSD), other mental health conditions, or altered resilience. Although certain behavioral and social factors have been associated resilience, little is known about the cellular and genomic mechanisms contributing to resilience or developing PTSD. Here, we present a novel model system called “Stress-in-a-dish” (SIAD) to study the molecular signature of chronic and acute stress in differentiated cortical organoids. Derived from human induced Pluripotent Stem Cells (iPSCs), mature organoids responded to cortisol with differential expression of genes. Many genes were associated with expected corticosteroid pathways, and some have not been previously linked to PTSD. A previously unknown potential contribution of astrocytes to the etiology of stress responses was also found. Our results suggest a novel paradigm for studying stress in a dish that opens up new opportunities to understand the biological basis of PTSD and resilience.

## Introduction

Studies have demonstrated that exposure to chronic stress and traumatic experiences can impact brain health and development^1^. For some individuals, such exposure may lead to Post-Traumatic Stress Disorder (PTSD), anxiety, or mood disorders^2^, while other individuals do not display adverse outcomes. This indicates that resilience, a measure of one’s susceptibility to handle stress without incurring mental health issues, is variable amongst individuals. However, little is known about the molecular and cellular factors that determine individual susceptibility to developing severe psychopathology after stress exposure and about the nature of a potential genetic basis underlying resilience.

Single or repeated exposure to stressors such as natural disasters including hurricanes, earthquakes and floods can leave people, particularly young people, struggling with anxiety, depression, poor sleep, and, in some cases, PTSD^3^. Current approaches to studying PTSD include genome-wide association studies (GWAS), animal studies, and post-mortem human brain tissue investigations as well studies which depend upon patients self-reporting PTSD symptoms^4-6^. A major difficulty with the diagnosis and treatment of PTSD is that there is an absence of objective biomarkers that can independently identify PTSD and reveal treatment efficacy. Our current methods also lack access to the molecular and genetic signatures of pre-mortem patient brains since–for obvious reasons–there is only so much (or so little) experimentation that can be done on living human brains.

Ideally, molecular studies would occur live in humans, comparing those that experience varying stress levels as well as among groups that identify as resilient or not. The obvious difficulty is accessing relevant sample material. However, the study of human brain tissues exposed to stressors is key to filling the gaps in knowledge at the genetic and molecular level that will help us fundamentally understand the expression of stress in the brain and how molecular interventions could improve resilience where it is lacking.

Since their invention, neural organoids have been shown to recapitulate many critical features of the cell-types and structures of the human brain cortex and are convenient for manipulation in genetic and molecular studies^7-10^. If generated from induced Pluripotent Stem Cell (iPSC) technology using skin or blood cells, one can cultivate organoids with the same genetic background of identified patients with neuropsychiatric disorders of interest. With those organoids, one can study the consequences of genetic variation on the execution of developmental programs of brain organogenesis as well as functional, chemical and epigenetic impacts.

A well known stress hormone is cortisol, which is released from the adrenal gland in stressful situations. Part of the response to cortisol release in the short term is increased glucose availability. However, long term exposure to cortisol can lead to negative health outcomes. It has been found that trauma can lead to the dysregulation of cortisol response. Patients reporting PTSD have altered cortisol levels depending on chronicity of symptoms^11^. Cortisol binds glucocorticoid receptors to transduce its signalling^12^. Downstream of cortisol, known activated pathways can include inflammatory and anti-inflammatory effects depending on length of exposure^13^. Prolonged cortisol exposure has also been shown to have neurotoxic effects in the hippocampus and cortex through changes in cell metabolism^14,15^. How cortisol is affecting resilience and how genetics can shape responses of neural pathways to cortisol is not sufficiently understood.

Here, we present a model system to study chronic and acute stress in an *in vitro* brain organoid system that is accessible to a wide range of genomic analyses. We confirmed responses of cortical organoids to a cortisol concentration mimicking physiologically high stress levels and performed single-cell RNA-seq to discover the differential responses of cortical cell types. We demonstrate SIAD as a new method to interrogate the contribution of stress on brain development and we expect that this model can prove a valuable tool to understand the etiology of PTSD, anxiety, and mood disorders.

## Results

### Validation of cortical organoids

As human fetal brain development is mostly inaccessible for longitudinal and molecular study, we hypothesized that we can recapitulate stress conditions by developing a cortical organoid system.

Human cortical organoids were differentiated from induced pluripotent stem cells (iPSCs) using previous protocols (Fig. 1A)^16^. At day 14, these organoids displayed neural rosettes as detected through Nestin and Foxg1 immunohistochemistry (Fig. 1B) and by day 50 express TUJ1 which indicate early neurons^17^. To further support the neural character, NeuN is assayed at day 75 and by day 100 and we observed strong staining for NeuO. To confirm that the organoids contained functional neurons, we conducted calcium imaging by applying Fluo4 indicator the organoids at day 150 which showed neuronal firing (Fig. 1B). To verify the cell subtypes contained within the human cortical organoids, we conducted single-cell RNA-seq on day 300 samples. All the major cell types of the human cortex including neurons, astrocyte progenitor cells, astrocytes, oligodendrocyte Precursor Cells (OPCs), and oligodendrocytes were present in our organoids (Fig. 1C, D), validating the recapitulation of *in vitro* human cortical brain tissue in a dish to study stress.

**Figure 1:**
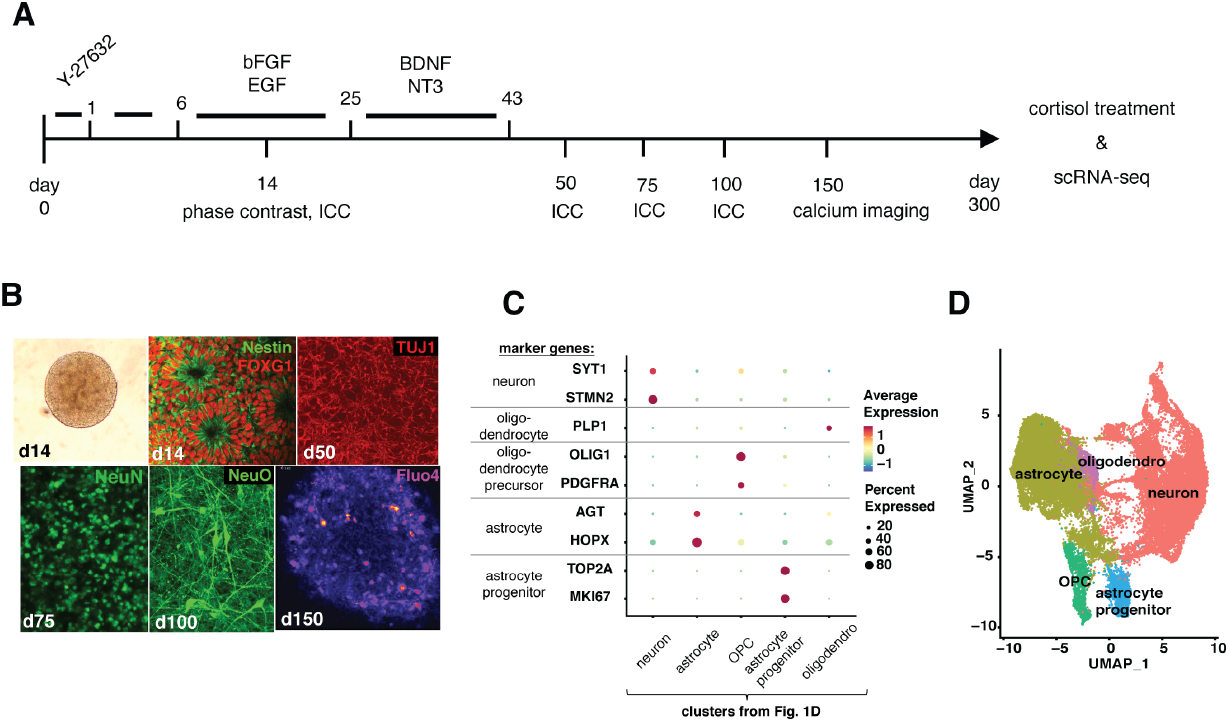
Human cortical organoids differentiated from induced pluripotent stem cells. A Timeline of cortical organoid differentiation and sampling timepoints (numbers = days). Top media additives, bottom assays being performed. B Phase contrast (d14), immunocytochemistry (d14, d50, d75, d100), and calcium imaging (d150) of human cortical organoids show presence of neural markers, neural morphology, and action potentials. C – Dot plot showing cortical cell types present in organoids determined through RNA expression analysis from Single Cell RNA-seq. Average expression of cell-type markers (Neuron – SYT1, STMN2; Oligodendrocytes PLP1; Oligodendrocyte Precursor Cell (OPC) OLIG1, PDGFRA; Astrocyte – AGT, HOPX and Astrocyte progenitor – TOP2A, MKI67) depicted by color with heatmap indicator. Size of circles indicate percentage of cells in the cell type class that express the marker. D – UMAP plot of first 2 principal components of clusters showing the major cell types found in day 300 cortical organoids. Colors depict markers of gene expression for major cortical cell types. OPC: oligodendrocyte progenitor cell, astro: astrocytes, astro.prog: astrocyte progenitor cell, oligodendro: oligodendrocytes.

### Cortical organoids respond to application of cortisol

The role of cortisol as a stress hormone is well established^18^. We devised a treatment schedule to mimic high stress by applying high, but still physiologically realistic, concentrations of cortisol. After washing out cortisol and returning to baseline for two days, a moderate to high pulse of cortisol was then applied. Organoids were subsequently dissociated and single-cell RNA-seq libraries prepared to measure the immediate cortisol effects on gene expression in human cortical cells (Fig. 2A).

**Figure 2:**
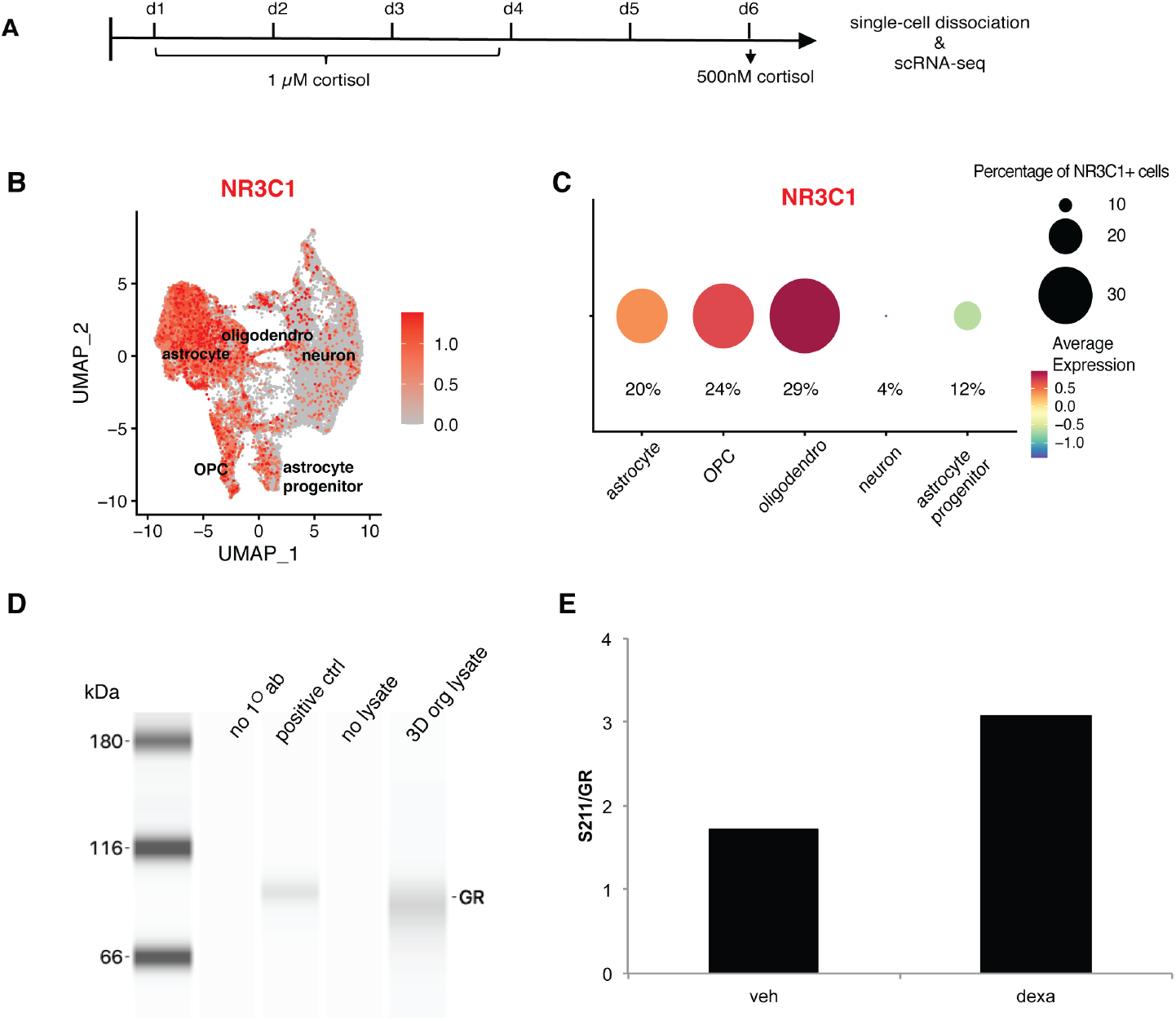
Stimulation of cortical organisms with cortisol. A - Timeline of cortisol treatment B – UMAP plot of cell-type clusters showing cell type expression of glucocorticoid receptor (GR) NR3C1. Red depicts normalized expression amount by cell type. C – Dot plot showing percentage of NR3C1 positive cells within each cortical cell type identified and their normalized expression (heatmap). D – Western blot analysis of presence of GR protein products in organoids. First lane: molecular weight marker, second lane: antibody-negative control, third label: U2OS cell lysate positive control, fourth lane: lysate-free negative control and fifth lane: organoid lysate. E – Bar chart showing ratio of S211 phosphorylated GR against total GR from quantitative western blot analysis of organoids with and without treatment with dexamethasone. OPC: oligodendrocyte progenitor cell, astro: astrocytes, astro.prog: astrocyte progenitor cell, oligodendro: oligodendrocytes.

Since stress responses to cortisol are mainly mediated through the glucocorticoid receptor (GR), we asked whether differentiated cells in the organoids expressed GR. By single-cell RNA-seq of cortical organoids, we found that 14.3% of the cells express the glucocorticoid receptor gene, *NR3C1* (Fig. 2B). In particular, a greater proportion of glial cells were NR3C1-positive compared to neurons (Fig. 2C). To confirm the presence of GR protein products, we performed western blotting with a GR antibody on organoid lysates (Fig. 2D). We detected a 94-97 kDa fragment that was a similar molecular weight to our positive control in U-2 OS cells and an absence of any bands in our lysate free or primary antibody negative controls (Fig. 2D).

Since cells in the organoid express GRs, we hypothesised that cortical organoids would respond to the direct application of cortisol. When cortisol binds to the glucocorticoid receptor, it becomes phosphorylated^19^. We thus tested for a response to cortisol in organoid cells by quantifying the phosphorylation of the glucocorticoid receptor by Western blotting. We applied 1µM cortisol to the organoids as it was a dose previously demonstrated to cause a robust response^15,20,21^, and collected the cells for protein analysis. For quantitative western blotting, we used an antibody specific to phosphorylated glucocorticoid receptor (p-GR). We observed 80% increase of p-GR signal relative to total GR protein in response to cortisol, confirming that they respond to cortisol application (Fig. 2E).

### Differential gene expression analysis reveals known and new stress response genes

Responses to stress can manifest as changes in gene expression^22^. Therefore, we asked whether cortisol treatment was able to elevate known genes involved in stress pathways in order to validate this new model for stress studies. To do this, single cell sequencing was conducted on mature d300 organoids treated with cortisol or control organoids to detect differential gene (DE) expression in response to cortisol.

Pseudobulk pathway analysis indicates significantly differentially regulated hypoxia pathway genes in response to cortisol (Fig. 3A). Eight genes out of 26 pseudobulk DE genes underlying this hypoxia signal are widely expressed in all cell types, but more strongly so in glia (Fig. 3B).

**Figure 3:**
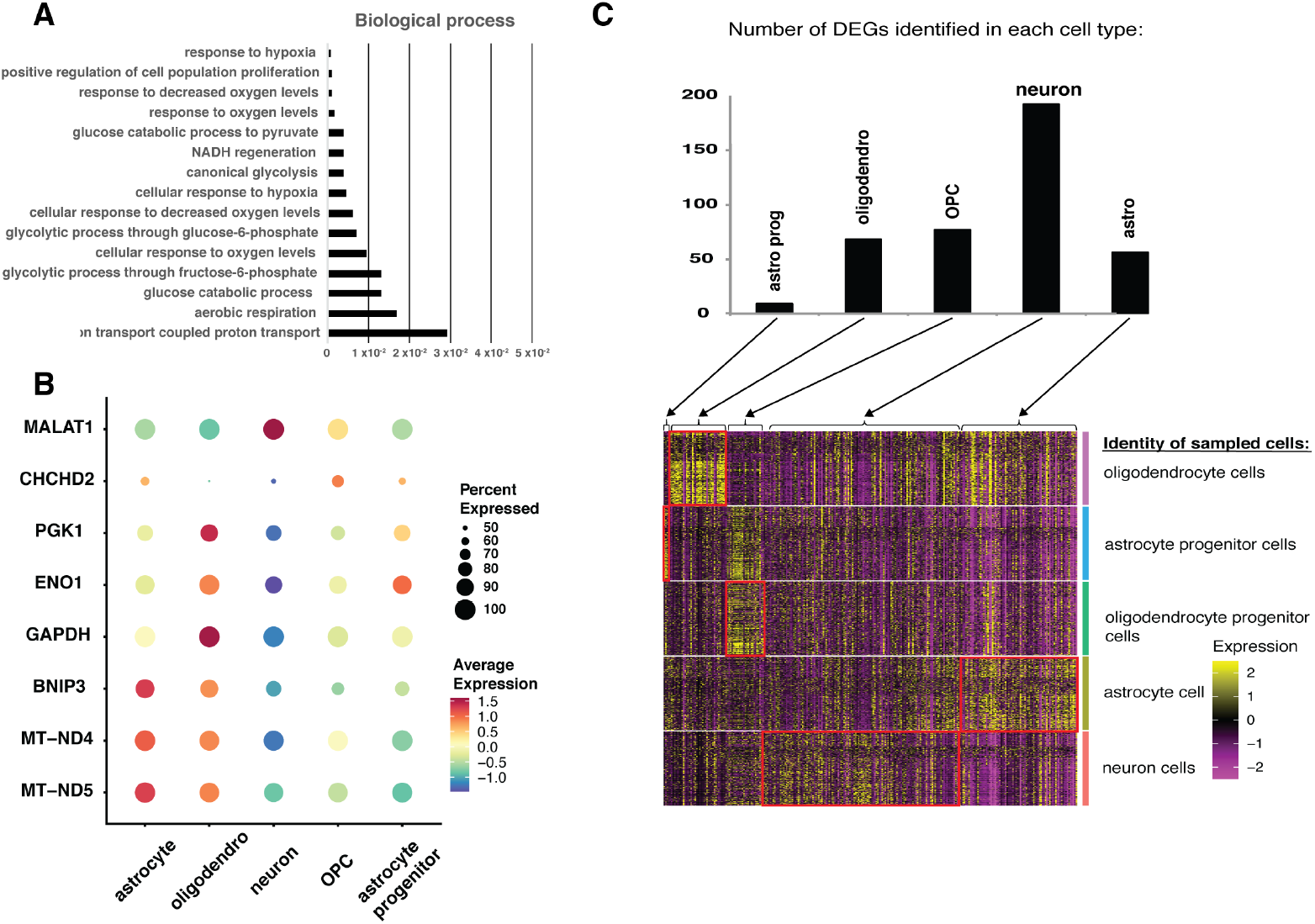
Differential gene expression analysis reveals known and new stress response genes. A- Bar graph showing Bonferroni-corrected p-value of pathways associated with differentially expressed genes B - Dot plot showing expression of hypoxia genes across the five major cell types C – Bar graph showing number of DE genes (top) and heat map (bottom) representation of expression of differentially expressed (DE) genes by cell type. Each cell-type specific group of rows represents 2000 randomly selected cells from an organoid. Each column represents DE genes grouped by cell types in which they were discovered. Heatmap: colors toward yellow is higher than expected expression and magenta is lower than expected. OPC: oligodendrocyte progenitor cell, astro: astrocytes, astro.prog: astrocyte progenitor cell, oligodendro: oligodendrocytes.

Along with expected differential expression of stress-related pathway genes, we grouped the responses by cell-types using known markers. (Fig. 3C). We observed a different number of DE genes when we performed the differential expression analysis in the five cell types separately. When looking where these cell-type candidate DE genes were expressed, we found two different patterns. For example, neuron DE genes did not appear to be neuron-specific. In contrast, astrocyte progenitor and oligodendrocyte DE genes were also only strongly expressed in these cell types (Fig. 3C).

### Cell-type specific gene expression reveals contribution of glial cells to stress response

We next asked whether there was overlap of DE genes between the cell types. The overlap analysis revealed very little overlap between the major cell subtypes (Fig. 4A, Supplementary Table 1). Most DE genes were only found in a single cell type. We quantified the similarities between the cell-type DE lists with a correlation matrix (Supplementary Fig. 1). The highest similarities were found between astrocytes, NR3C1+ astrocytes, and NR3C1-positive cells.

**Figure 4:**
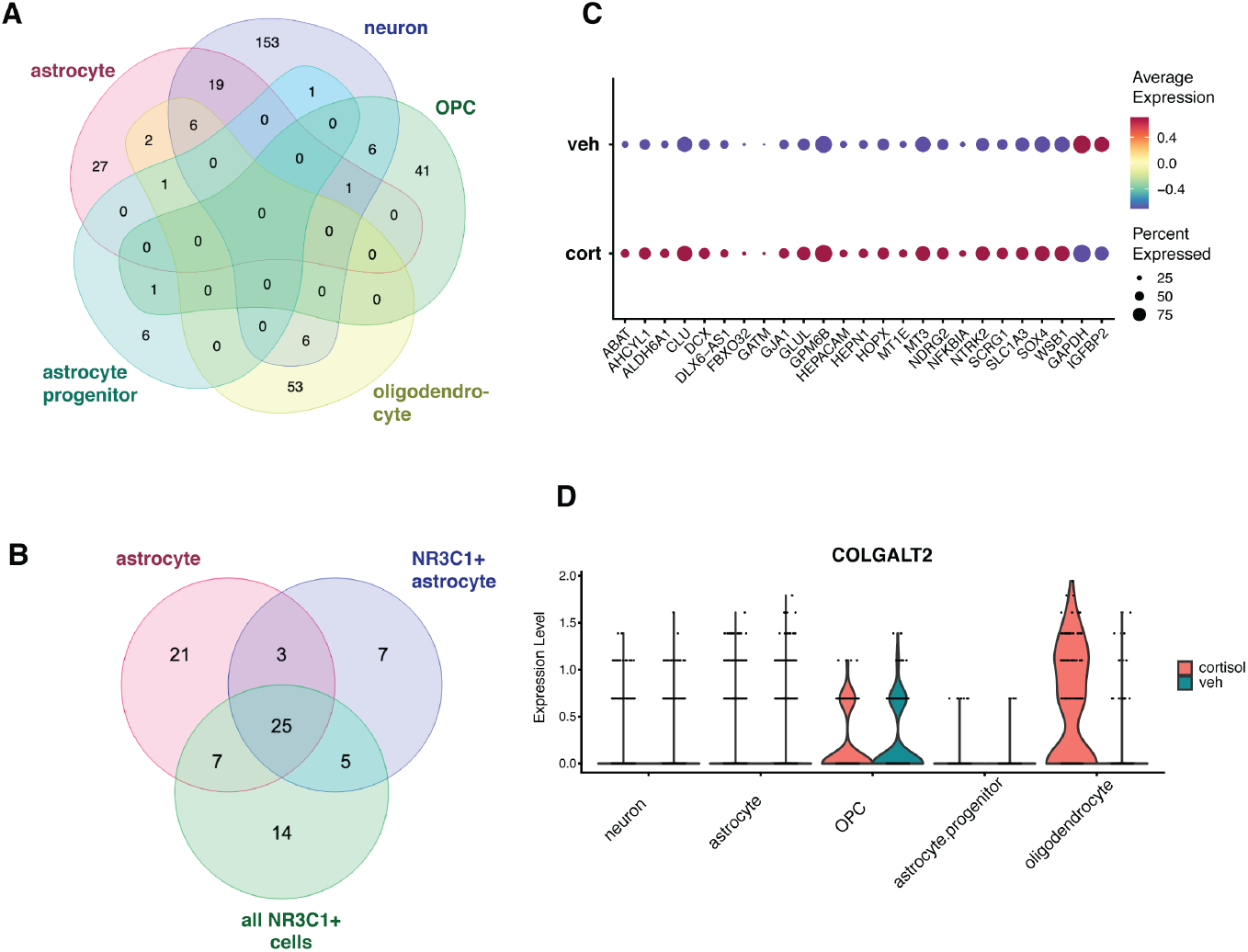
Cell-type specific gene expression reveals contribution of glial cells to stress response. A - Venn diagram of overlap between lists of DE genes by cortical cell types. B - Venn diagram of overlap of DE lists from astrocytes (red), NR3C1 positive astrocytes (blue) and all NR3C1 positive cells (green). C – Dot plot showing expression levels of 25 overlapped genes in the cortisol-treated (cort, bottom) versus vehicle treated (veh, top) organoids. Size of circles indicated percentage of cells expressing the gene of interest. Color of circle depicts normalized expression level. D - Violinplot of COLGALT2 expression across five major cell types. Red: cortisol treated sample, green: vehicle-treated samples. OPC: oligodendrocyte progenitor cell, astro: astrocytes, astro.prog: astrocyte progenitor cell, oligodendro: oligodendrocytes.

A Venn diagram between these three groups indicates that 28 of the 56 DE genes in astrocytic populations are driven by direct NR3C1+ expression. The expression of these genes and the percentage of the subtype that express the DE genes suggest that astrocytic gene expression is a major component of the response to cortisol (Fig. 4B).

A core set of 25 genes is shared between all three lists. 23 out of the 25 genes are upregulated in response to cortisol (Fig. 4C). Most genes have support in the literature for a role in cortisol or stress responses and have been associated in GWAS to PTSD and resilience (see Discussion). However, comparing our findings to a list of genes where SNPs have been associated with PTSD, significant overlap has not appeared (Supplementary Table 2)^4^. Interestingly, a compiled list of genes revealed through functional genomics across human and animal tissues do show strong enrichment for astrocye, NR3C1+ all, NR3C1+ astro, as well as 25 core genes (Supplementary Table 3)^5^.

To further validate this model of stress, we conducted a pathway analysis in an unbiased fashion on whether known and unknown cellular and genetic pathways related to PTSD were differentially regulated in response to cortisol stimulation (Supplementary Table 4). We find pathway analysis points to the corticosterone pathway gene list being significantly enriched (p < 0.00001).

In addition to finding sets of genes enriched in response to cortisol, we have also identified novel genes that are significantly differentially expressed but are not traditionally related to stress responses, such as ALDH6A1 and COLGALT2. ALDH6A1, one of the 25 core genes (Fig. 4C) encodes an aldehyde dehydrogenase. Mutations in ALDH6A1 have been associated with changes in myelin in the brain and thinning of the corpus callosum^23^. COLGALT2 (Fig. 4D) encodes the procollagen galactosyltransferase 2 which glycosylates hydroxylysine residues in collagen^24^. Overall, our results suggest that SIAD is a powerful novel method to study molecular and genetic mechanisms of stress responses without requiring live patient experimentation.

## Discussion

In this study, we introduce a new Stress-In-A-Dish (SIAD) system to model the impact on the molecular level of stress on the development and functionality of the human brain. By recapitulating stress conditions “in a dish”, this approach is expected to greatly increase the potential of studying and diagnosing resulting mood disorders, ultimately leading to development of treatments for patients in a non-invasive, rapid and personalized manner.

### SIAD is a validated, unbiased model of stress

With an unbiased survey of genome-wide expression on organoid cells, SIAD has not only shown a molecular response to cortisol that one might expect from a literature search but has also revealed novel potential candidate genes for stress responses. Organoids express the NR3C1 glucocorticoid receptor gene as expected and gene products significantly associated with corticosterone drug interaction. Two further, relevant gene lists from the literature include the PTSD GWAS candidate genes from Nievergelt et al. and candidate genes from Stankiewicz et al., that were collated from functional genomics studies (3 human post-mortem brain transcriptomics and 79 animal studies). When comparing our DEG lists to Stankiewicz and Nievergelt we find significant overlap with the Stankiewicz list, but not with the Nievergelt list (Supplementary Tables 2 and 3). Interestingly, the Stankiewicz and Nievergelt lists are distinct and show no overlap apart from one gene (SEMA3B). This may explain why we find significant association only with the Stankiewicz list and not with Nievergelt list.

When taking a wider lens and asking which genes are examinable in our proposed model, the overlap becomes even larger - 88% of Stankiewicz genes are expressed and present in our organoid model, compared to 60% of Nievergelt genes. One might speculate that such divergence in coverage indicates the difficulty linking of GWAS-generated SNPs to candidate genes, and perhaps it is to be expected that gene lists from our own functional model more closely resemble findings from other functional studies, such as Stankiewicz et al. Importantly, we matched candidate genes from human post-mortem brain and animal studies indicating that our SIAD model has the yield and efficacy of more intensive functional studies.

### Novel cell types found to potentially transduce stress responses

Our unbiased assay allowed us to see glial cells as important transducers of stress hormones in the brain. This demonstrates the importance and value of growing older organoids, validating the presence of glia in organoids and allowing enough time for the maturation of astrocytes and oligodendrocytes. While future studies will investigate the role of glial transduction in stress responses, it is interesting to note that white matter reduction is common in children with adverse childhood events^25^.

Unlike non-guided organoids that were stimulated with the synthetic cortisol-like steroid dexamethasone^26^, it could be that by virtue of being younger abrogated the identification of the impact of glia in those organoids, and revealing instead progenitor cell effects. It might be concluded that organoids that best recapitulate the mature human brain cell population and connectivity will be best placed to study brain disorders.

### Involvement of hypoxia responses to cortisol treatment

Surprisingly, we found an enrichment of hypoxia-related genes in the cortisol response of organoids. These changes appear to be localised to the glial cell population. Importantly, these results are important with respect to the potential impact of neonatal hypoxia and ischemia to the development of stress-related responses in adulthood^27^. A tempting hypothesis to test is whether hypoxia can predispose children to mis-regulated stress responses in adulthood. What is the impact of maternal prenatal stress on the developing child?

### Discovery of new candidate genes

A gene that neither appears on the functional genomics (Stankiewicz et al.) nor GWAS list (Nievergelt et al.), but which was found to be upregulated in response to cortisol exposure, is COLGALT2, a galactosyltransferase which glycosylates hydroxylysine residues in collagen^24^. O-glycosylation of collagen is important for the correct collagen assembly and deposition in the extracellular matrix (ECM)^28^.

A recent multi-omic study of post-mortem brain and blood plasma in patients with a diagnosis of PTSD or Major Depressive Disorder identified COL4A1 as a top candidate gene and more broadly ECM structural components as a top candidate pathway^6^.. Interestingly, unlike its isoenzyme COLGALT1, which shows a wide expression across tissues, COLGALT2 seems to be specifically expressed in brain tissues, with highest expression in the spinal cord and corpus callosum^24^. Single-nuclei RNA-seq data shows that COLGALT2 is most abundantly expressed in oligodendrocytes (Supplementary Figure 2).

Emerging evidence suggests that extracellular matrix (ECM) alterations can occur with stress^29^. In a murine animal model of stress, it was observed that in the Ventral Tegmental Area (VTA) of male mice Colgalt2 gene expression was upregulated^30^.

In addition to the link between stress and the extracellular matrix, both are also associated with hypoxia, captured by the pathway analysis of DE genes from all cell types (Fig. 3A). Hypoxia inducible factors (HIF) have been shown to regulate ECM synthesis and remodeling^31^. For example, cortical organoids cultured for 48 hours under hypoxia showed downregulation of COL4A6, a differentially expressed gene in zebrafish model of chronic unpredictable stress^32^.

In conclusion, COLGALT2 and the other novel DE genes are interesting avenues to follow up in terms of understanding the transduction of the stress response in the brain. For their study, SIAD is an accessible model, for example, to study ECM and collagen formation and modification in stress.

### The future of SIAD models

It is clear that organoids do not fully model the whole brain, and with it the complex inter-regional communication including the limbic system that are known to be intimately involved in stress disorders such as PTSD. Here, current attempts to create multi-regional organoids could improve the modelling of such interactions^33^. That said, organoids can be expected to bridge important knowledge gaps to initially identify disorders, mechanisms and the effects of underlying genetic variants and treatment prior to entering drug candidates to expensive clinical trials. Positive and negative interactions between drug candidates and genotypes can be elucidated to filter targets and de-risk clinical trial efforts. SIAD is a flexible model as iPSC cell lines from patients with low or high resilience to stress can be studied to elucidate mechanisms of resilience that could be targeted pharmacologically.

## Acknowledgements

We would like to thank members of the Urban and Carrion labs for useful discussions and revising the manuscript. We thank Agnieszka Kalinowski for useful discussions. We thank the Beckman Center’s Cell Sciences Imaging Facility (CSIF) for assistance with microscopy, Stanford Genetics Bioinformatics Service Center and Stanford Research Computing Center for providing computing resources. This work was funded by the National Institutes of Health and a generous gift from Bruce Blackie.

---

**Supplementary Figure 1:**
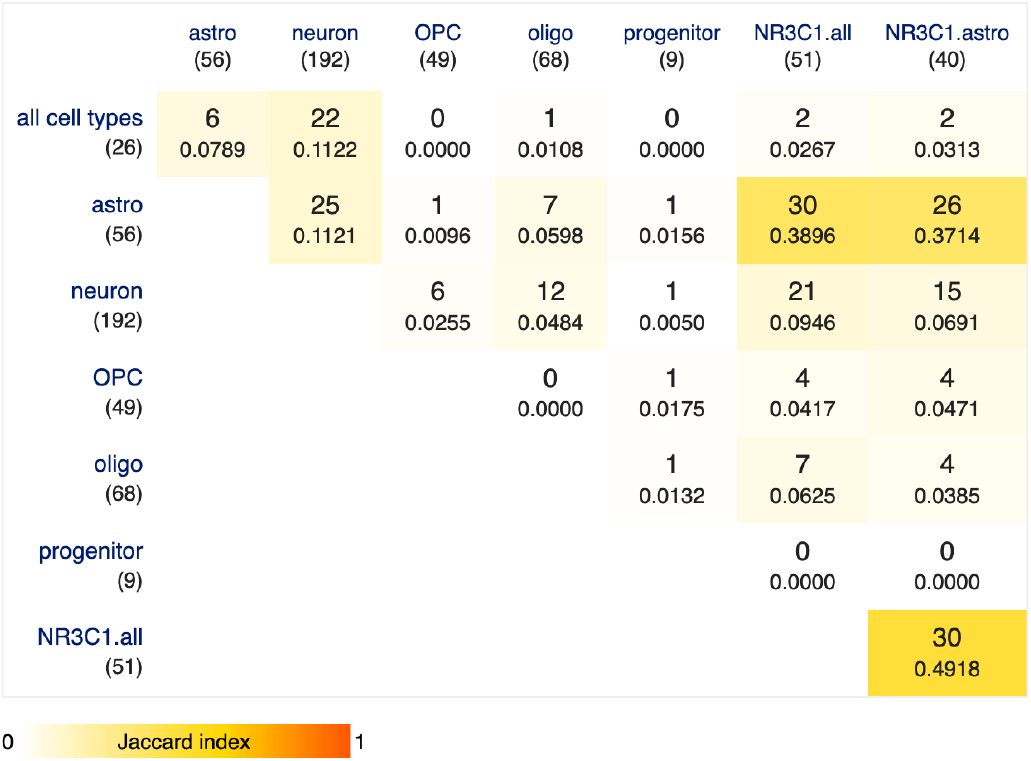
Correlation matrix of similarities between DE lists Integer values represent number of genes that overlap and decimal numbers are values of the Jaccard Index

**Supplementary Figure 2:**
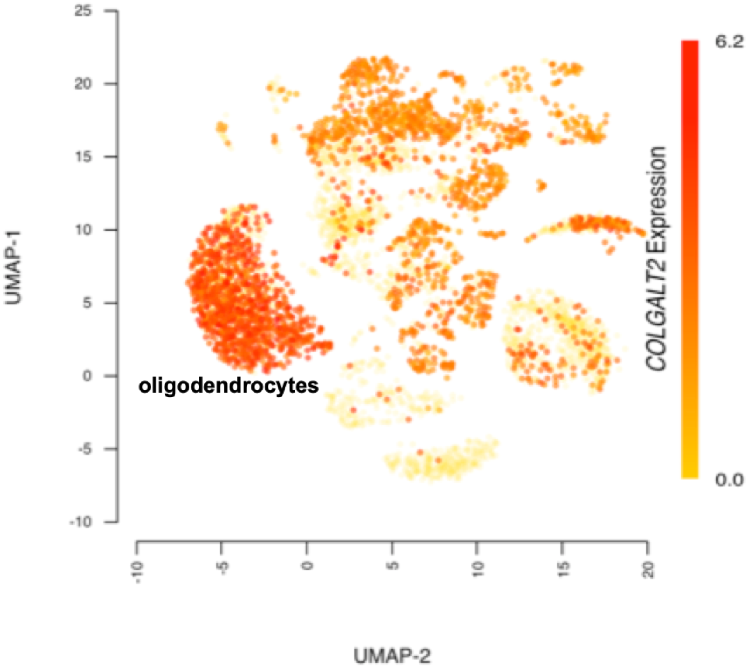
Cell-type specific gene expression of COLGALT2 in human post-mortem brain. UMAP plot of single-nuclei gene expression of COLGALT2 in dorsolateral prefrontal cortex. Data taken from PsychSCREEN, MultiomeBrain-DLPFC.

## Materials & Methods

### hiPSC: donors, reprogramming and maintenance

Human induced pluripotent stem cells (iPSCs) were reprogrammed from human dermal fibroblasts (HDFs) using Sendai’s CytoTune-iPS 2.0 Sendai reprogramming kit (Invitrogen, A16517). HDF cell lines were established from skin biopsies collected at Stanford from two healthy female donors in compliance with IRB (insert ethics protocol). iPSCs were maintained on matrigel with mTesr medium (Manufacturer) until differentiation. Pluripotency was verified by staining for Tra1-60, Nanog and SSEA-4 with standard immunocytochemistry as described previously (urban ref).

### Cortical organoids

Cortical brain organoids prepared as described in Pasca et al., Nat Methods 12, 671–678 (2015), with the following modifications. iPSC colonies were not lifted as a whole to form embryoid bodies of varying sizes. Instead iPSCs were dissociated into cell solution and uniformly sized embryoid bodies were prepared using AggreWell Microwell plates (STEMCELL Technologies Catalog #34811).

### ICC

Human cortical organoids were fixed for 20 minutes in 4% paraformaldehyde (PFA) diluted in phosphate-buffered saline (PBS) at room temperature. The fixed organoids were then washed three times for 10 minutes each with PBS to remove residual PFA. For permeabilization and blocking, the organoids were incubated in a solution of 0.3% Triton X-100 and 5% normal goat serum (or other relevant serum) in PBS for 1 hour at room temperature.

Following blocking, the organoids were incubated with primary antibodies diluted in the blocking solution at 4° C overnight on a rocker. After primary antibody incubation, the organoids were washed three times for 10 minutes each with PBS.

Secondary antibodies, diluted in blocking solution, were applied for 2 hours at room temperature in the dark. The secondary antibodies used were: goat anti-mouse Alexa Fluor 488, goat anti-rabbit Alexa Fluor 594, and goat anti-guinea pig Alexa Fluor 647 (all at 1:1000 dilution, Invitrogen). After secondary antibody incubation, the organoids were washed three times for 10 minutes each with PBS.

The stained organoids were imaged using a confocal microscope (e.g., Zeiss LSM 880). Z-stack images were acquired to capture the three-dimensional structure of the organoids. The images were processed and analyzed using Fiji.

### Calcium imaging

Human cortical organoids were cultured and maintained in a defined media in an incubator at 37° C with 5% CO_2_. For calcium imaging, organoids were incubated with 5 µM of the cell-permeant calcium indicator Fluo-4 AM (Invitrogen) and 0.02% Pluronic F-127 for 45 minutes at 37° C in the dark. After incubation, the loading solution was gently replaced with fresh HBSS, and the organoids were allowed to de-esterify the dye for an additional 15 minutes at room temperature.

Imaging was performed on an DeltaVision OMX microscope. Fluo-4 was excited using a 488 nm laser, and emission was collected at 515-545 nm. Time-lapse movies were acquired at a frame rate of 1 Hz for a duration of up to 10 minutes. Acquired videos were analyzed using Fiji image processing software.

### Glucocorticoid treatment

Human cortical organoids were maintained in culture in a defined neural basal media supplemented with B27, N2, and other necessary growth factors. Before treatment, the organoids were transferred to fresh media to ensure a clean baseline. The culture media was then supplemented with cortisol at a final concentration of 1 µM. The organoids were treated for a period of 72 hours in a humidified incubator at 37 degrees Celsius with 5% carbon dioxide. A control group was run in parallel, receiving only the vehicle used to dissolve the cortisol, to account for any effects of the solvent itself.

Following the 72-hour treatment period, the organoids were allowed grow in in their normal growth medium for two days again before being treated with 500 nM cortisol to simulate a trigger. In order to model the immediate gene expression response to the trigger, organoids were harvested after one hour and immediately processed for downstream analysis.

### Western Blot

Protein samples were prepared for analysis with a WES machine (ProteinSimple, San Jose, CA). After lysing cells in RIPA buffer, protein sample concentrations were determined using a BCA protein assay kit (Pierce, Rockford, IL) according to the manufacturer’s instructions. Samples were then diluted to a final concentration of 1 mg/mL in a master mix containing 0.4× sample buffer, 1× fluorescent master mix, and 2× dithiothreitol (DTT). The mixture was denatured by heating at 95 degrees Celsius for 5 minutes.

For each sample, a WES 12-230 kDa Separation Module (ProteinSimple) was used. The module consists of a pre-filled plate with reagents and capillaries. Each capillary was loaded with a blocking reagent, a stacking and separation matrix, and the protein sample. The instrument performed automated electrophoresis, separating proteins by size. Separated proteins were then immobilized on the capillary walls via a proprietary photoactivated capture chemistry.

Following immobilization, the capillaries were washed, and the target protein was probed. The primary antibody was diluted in Antibody Diluent 2 and incubated for 30 minutes, followed by a wash. A horseradish peroxidase (HRP)-conjugated secondary antibody was then added and incubated for 30 minutes. The capillaries were then washed to remove any unbound secondary antibody. Chemiluminescent substrate was then added for signal detection.

Data was analyzed using the Compass for Simple Western software (ProteinSimple). The software automatically generated a virtual blot and an electropherogram for each sample. The area under each protein peak in the electropherogram was used for quantitative analysis of protein expression. The signal was normalized to total protein using the built-in normalization feature of the software.

### Single-cell sequencing

On day 300 of differentiation, organoids were dissociated as previously described in Sloan et al., Neuron 95, 779-790 (2017). Single-cell RNA-seq (scRNA-seq) libraries were prepared using 10x Genomics Chromium Single Cell 3’ Reagents v3 following manufacturer’s instructions. Briefly, the concentration of single cells in solution was determined using Trypan Blue staining. The cell solution was loaded onto a Chromium Chip B to capture three to four thousand cells in droplets containing the reverse transcription reagents. After reverse transcription, the now barcoded cDNA was recovered and amplified for 12 PCR cycles. After qualitative and quantitative control of the cDNA, the final libraries were constructed by fragmenting the cDNA, End Repair, and A-Tailing. After adapter ligation, the libraries were amplified for 11 PCR cycles. The libraries were sequenced on a Illumina NovaSeq machine to achieve a sequencing depth of about 80,000 reads per cell.

Single-cell RNA-seq (scRNA-seq) data were analyzed using the Seurat package (version 5.0) in R (version 4.2.0). Raw gene-by-cell count matrices, generated from the 10x Genomics Cell Ranger pipeline, were imported into R using the Read10X function.

Each dataset was first filtered to remove low-quality cells and potential doublets. The remaining high-quality cells were then normalized using the SCTransform function, follwed by RunPCA, RunUMAP, FindNeighbors, and FindClusters.

Cell clusters were annotated based on the expression of canonical marker genes identified from existing literature. Differentially expressed genes for each cluster were determined by comparing the gene expression of cells within a cluster to those in all other clusters using the FindAllMarkers function. Genes with an adjusted p-value of less than 0.05 were considered significant.

